# Haplotypes variations of yellow stripe like (*TaYSL*) genes are associated with grain iron and zinc contents in wheat (*Triticum aestivum L.*)

**DOI:** 10.64898/2026.06.17.732851

**Authors:** Kainat Abbasi, Humaira Qayyum, Samar Naseer, Mengjin Sun, Muhammad Adeel Quraishi, Youshaa Danyal, Yuanfeng Hao, Zhonghu He, Awais Rasheed

## Abstract

The availability of pangenome and resequencing of wheat collections have facilitated the discovery of gene-trait associations in wheat. Yellow stripe-like (YSL) proteins play a key role in the uptake and translocation of metals and yet have not been fully identified and analyzed at the genome-wide level in wheat. In this study, 26 *TaYSL* genes were identified and divided into four distinct clades, each clade sharing similar domains and motif compositions. Most genes were upregulated under iron deficiency, whereas homoeologs of *TaYSL1* were downregulated. Both SNP-based and haplotype-based association studies were used to dissect the role of TaYSLs underpinning grain iron contents (GFeC) and zinc contents (GZnC) in wheat. *TaYSL6-2B* and *TaYSL16-1A* haplotypes showed strong association with GFeC, and *TaYSL14-6A* showed strong association with GZnC in multiple field trials. The distribution of favorable haplotypes in global wheat collection of ∼3000 accessions showed that majority of haplotypes were more prevalent in landraces and winter wheat compared to modern cultivars and spring types, indicating their potential for use in breeding. The combination of favorable haplotypes of three YSL genes associated with GFeC and GZnC were very rare, and most of the wheat accessions has single or double favorable haplotypes. These findings provide the first comprehensive characterization of the *TaYSL* gene family in wheat and identify significant SNPs and elite haplotypes that can be utilized for genetic improvement and biofortification.

## Introduction

Wheat (*Triticum aestivum* L.), one of the most important staple food crops in the world, contributes more than 50% of the diet and up to 60% of daily intakes of iron (Fe) and zinc (Zn) in several developing countries (Cakmak et al., 2010). Fe is indispensable for the survival of living organisms, as it is integral to key physiological processes including citric acid cycle, electron transfer in the respiratory chain, biological nitrogen fixation, protein and nucleic acid synthesis (Davila-Hicks et al., 2004). Despite being the fourth most abundant element in the earth’s crust, low Fe bioavailability in aerobic, calcareous and high pH soils results in chlorosis primarily due to impaired chlorophyll production (Buckhout et al., 2009; Morrissey et al., 2009; Barker and Stratton, 2015). In humans, Fe deficiency affects nearly two billion people globally, causing anemia, impaired cognitive development, and compromised immunity, with highest risk among children and pregnant women in developing countries (Vasconcelos et al., 2017; Lopez et al., 2016). Thus, enriching crops with micronutrients through biofortification presents a sustainable solution to improve nutritional security (Blancquaert et al., 2017). Therefore, it is important to discover genes involved in Fe homeostasis, and biofortification to use this knowledge in wheat breeding.

Plants have evolved two primary Fe uptake mechanisms, which differ among species (Kobayashi et al., 2012). Non-graminaceous plants acquire Fe through the reduction-based strategy (Strategy I), in which H -ATPases acidify the rhizosphere, FRO2 reduces Fe³ to Fe² , and IRT1 transports Fe² into root epidermal cells (Santi et al., 2009; Waters et al., 2002; Vert et al., 2002). Graminaceous plants utilize chelation-based strategy (Strategy II), where roots secrete mugineic acid-type phytosiderophores (PS) via TOM transporters to chelate Fe, and the resulting Fe³ -PS complexes are then taken up by Yellow Stripe1 (YS1) or YS1-like (YSL) transporters (Nozoye et al., 2011; Yordem et al., 2011).

YSL proteins, members of Oligopeptide Transporter (OPT) family, play a vital role for metal homeostasis by facilitating long-distance transport of metal-chelates (Feng et al., 2017; Ishimaru et al., 2010). The *YSL* gene derives its name from the “yellow stripe” phenotype observed in the *zmys1* mutant, which exhibits interveinal chlorosis due to a defect in Fe acquisition (Curie et al., 2001). With the sequencing of multiple plant genomes, it has become evident that higher plants contain four distinct and well-conserved groups of YSL proteins, one is specific to grass species (Yordem et al., 2011). These proteins are identified by highly conserved OPT domain which contains a conserved sequence K(F/R)L(T/P/A)(Y/F)PSG(T/L/S)ATA(V/M/H)LIN (Gendre et al., 2007).

To date, multiple YSL family members have been characterized in model plants. These include eight members in the dicot *Arabidopsis thaliana* (DiDonato Jr et al., 2004), 18 members in *Oryza sativa* and *Zea mays* (Zhang et al., 2018; Song et al., 2024), 19 in *Brachypodium distachyon* (Yordem et al., 2011) and 26 members in *Triticum aestivum* (Kumar et al., 2019). In rice, *OsYSL2* facilitates the uptake of metal-NA complexes to shoot and seeds while *OsYSL6* functions to prevent metal hyperaccumulation and mitigate Mn toxicity (Koike et al., 2004; Sasaki et al., 2011). *OsYSL9* is responsible for transporting Fe^2+^-NA and Fe^3+^- DMA from the endosperm to the developing embryo (Senoura et al., 2017). *OsYSL15* mediates phloem dependent Fe unloading and subsequent loading into seeds (Inoue et al., 2009). Furthermore, *OsYSL16* absorbs Fe^3+^-DMA and distributes Fe^3+^-DMA from the xylem to neighboring cells and *OsYSL18* has role in Fe^3+^-PS complexes uptake, mediates xylem-phloem Fe translocation (Kakei et al., 2012; Aoyama et al., 2009). In wheat, *TaYS1A* is involved in providing resistance against yellow rust by modulating SA-induced signaling (Islam et al., 2020). Similarly, *TaYSL15-6B* transports Cd-NA from roots, prevents Cd accumulation in the grains (Yao et al., 2025).

In this study, we identified *YSL* family genes in wheat genome, investigated tissue specific expression of *TaYSL* genes and their response to Fe deficiency, identified the genetic variants of *YSL* genes associated with grain Fe and Zn concentrations, and distribution of elite haplotypes in global wheat collection. This research provides basis for understanding the evolution and putative functions of *TaYSL* family genes in mineral biofortification.

### Materials and methods Germplasm and phenotyping

We used a diversity panel consisting of 145 landmark cultivars (Hao et al., 2020). The list of cultivars is presented as Supplementary Table S1. The diversity panel was phenotyped for grain iron contents (GFeC) and grain zinc contents (GZnC) using EDXRF across three growing seasons (2021-2023) at Gaoyi (GY) and Gaocheng (GC) in China (Sun et al., 2023; Naseer et al., 2024). A spring wheat cultivar, Zincol-16, was used and is notable for its high GFeC and GZnC with an average of 52 mg kg^−1^ and 48 mg kg^−1^.

### Characterization of TaYSL proteins in wheat

To identify TaYSL proteins in wheat, Hidden Markov Model (HMM) profile for the OPT domain (PF03169) was used to scan the wheat proteome using HMMER 3.0 (E value < 1×10^−5^) (Finn et al., 2014; Eddy et al., 2011). Sequences were filtered to retain only those containing complete domain and YSL signature sequence (Feng et al., 2022). The putative TaYSL proteins were further validated using the NCBI Conserved Domains Database (https://www.ncbi.nlm.nih.gov/Structure/) and the Simple Modular Architecture Research Tool (SMART) program (https://smart.embl.de/) (Marchler-Bauer et al., 2015; Letunic et al., 2021). Physicochemical properties of TaYSL proteins were analyzed using ExPASy (https://web.expasy.org/protparam/) , subcellular localization was predicted with WoLF PSORT (https://wolfpsort.hgc.jp/), and transmembrane helices (https://dtu.biolib.com/DeepTMHMM) were identified using DeepTMHMM (Duvaud et al., 2021).

The phylogenetic tree was constructed tree using neighbor-joining (NJ) method with 1000 bootstrap replicates (Tamura et al., 2021) which was visualized through Interactive Tree of Life (iTOL) (https://itol.embl.de/). The *TaYSLs* gene structure were mapped using the Gene Structure Display Server (GSDS) tool (http://gsds.cbi.pku.edu.cn/) (Hu et al., 2015). Conserved motifs were determined with Multiple Expectation Maximization for Motif Elicitation (MEME) (https://memesuite.org/meme/tools/meme), and data visualization was conducted with TBtools (Bailey et al., 2015; Chen et al., 2020). Cis-regulatory elements were characterized in the promoter sequences using the PlantCARE database (http://bioinformatics.psb.ugent.be/webtools/plantcare/html/) (Lescot et al., 2002).

### Expression pattern analysis of *TaYSL* family genes

We obtained tissue-specific *TaYSL* genes RNA-seq data of the wheat cultivar Chinese Spring encompassing various developmental stages from the WheatOmics 1.0 (http://202.194.139.32/expression/wheat.html) in Transcripts Per Kilobase Million (TPM) (Ma et al., 2021). Using transcriptomic data from our study (Qayyum et al., 2026), we extracted TPM values for *TaYSL* genes from both control and iron-stressed wheat root and leaf samples to analyze their expression under Fe deficiency conditions. Expression patterns were visualized using TBtools, and heatmaps were generated based on log_2_(TPM + 1) values (Chen et al., 2018). Similarly, the expression profiles of *TaYSL* genes were obtained from the published transcriptome dataset (Guo et al., 2025). The data was used to analyze gene expression patterns during seed development, particularly in the endosperm and embryo.

### Plant cultivation and growth parameters

Seeds of the wheat cultivar Zincol-16 were surface-sterilized prior to germination on moist filter paper under dark conditions. Seedlings were grown hydroponically in a controlled greenhouse under a 16/8 h light/dark cycle at 25°C using full-strength Hoagland’s solution (pH 5.8) which includes macronutrients (1mL/L 1M KH_2_PO_4_, 1mL/L 2M MgSO_4_.7H_2_O, 2.5mL/L 2M Ca(O_3_)_2_.4H_2_O, 2.5mL/L 2M KNO_3_), 1mL/L micronutrient stock solution (H_3_BO_3_, MnCl_2_.4H_2_O, ZnSO_4_.7H_2_O, CuSO_4_.5H_2_O, H_2_MoO_4_.H_2_O) and 1.5mL/L Fe (III)- EDTA. To induce Fe stress, one-week-old seedlings grown in full nutrient solution were transferred to a half-strength Hoagland solution without Fe (III)-EDTA salt. Root and leaf samples were harvested on 7, 14, and 21-days, snap-frozen and stored at -80°C.

Total RNA was purified from the collected samples using the TRI Agent (Invitrogen, USA) Extraction Kit, while NanoDrop 2000 spectrophotometer measured RNA concentration. Complementary DNA (cDNA) was synthesized from RNA using the ABScript III RT Master Mix (ABClonal). The Genious 2x SYBR green rapid qPCR mix (ABclonal) amplified target sequences in real-time with *TaActin* providing internal reference for relative quantification. Gene-specific primers for qRT-PCR were designed with NCBI Primer-BLAST (Supplementary Table S2). Relative *TaYSL* gene expression values were computed using 2^−ΔΔCT^ method (Schmittgen and Livak, 2008) from three independent biological replicates.

### Data Analysis

The whole genome resequencing data was obtained from the Wheat Union database (http://wheat.cau.edu.cn/WheatUnion/c_18/). The variants were filtered by applying a minor allele frequency (MAF) greater than 5% and a maximum missing rate threshold of 0.2. TASSEL software (version 5.0) was used to perform a general linear model (GLM)-GWAS at p=0.01 using the values of GFeC and GZnC in six environments (21GC, 21GY, 22GC, 22GY, 23GC, 23GY) including average (Bradbury et al., 2007). Haplotypes were constructed using RTM-GWAS software v1.2, which is publicly available at https://gitee.com/njau-sri/rtm-gwas (He et al., 2017). The output vcf from RTM-GWAS was used as a marker dataset for association analysis in TASSEL. Linkage disequilibrium (LD) heatmaps of significant blocks were generated using LDBlockShow (https://github.com/BGI-shenzhen/LDBlockShow) (Dong et al., 2021). The geographical distribution of the significant haplotypes was studied using 2789 accessions from the Wheat Union Database, which have well-documented information on status (landrace/cultivar), growth habit (spring/winter), country, and continent of each accession.

## Results

### Identification and characterization of TaYSL members in wheat

In this study, HMM searches identified 67 non-redundant TaYSL proteins. Our analysis using the IWGSC RefSeq v1.1 assembly led to a revised list of TaYSL family members. We found that previously *TaYSL7-7D* and *TaYSL7-7A2* corresponded to identical sequences while *TaYSL23-5A*, *TaYSL3-2D*, *TaYSL4-2A*, *TaYSL16-1D*, and *TaYSL19-2B* had incomplete domains and were removed (Kumar et al., 2019). Additionally, *TaYSL23-5A* was reassigned to a newly identified gene with a complete OPT domain. The detailed information of these proteins including amino acids, exons, introns, subcellular localization, isoelectric point, molecular weight, instability index, aliphatic index, grand average of hydropathy, and the number of transmembrane helices is listed in Supplementary Table S3.

To explore the evolutionary relationship, a phylogenetic tree was constructed using aligned amino acid sequences from TaYSLs along with previously reported OsYSLs, AtYSLs, and BdYSLs (Figure 1). Based on sequence similarities, YSLs were classified into four distinct groups. Group I and III were the largest subgroups containing 33 members, group IV contained 28 YSLs, while group II comprised 19 members. Notably, group IV comprised monocot YSLs, whereas the remaining subgroups included both monocot and dicot members.

**Figure 1.**
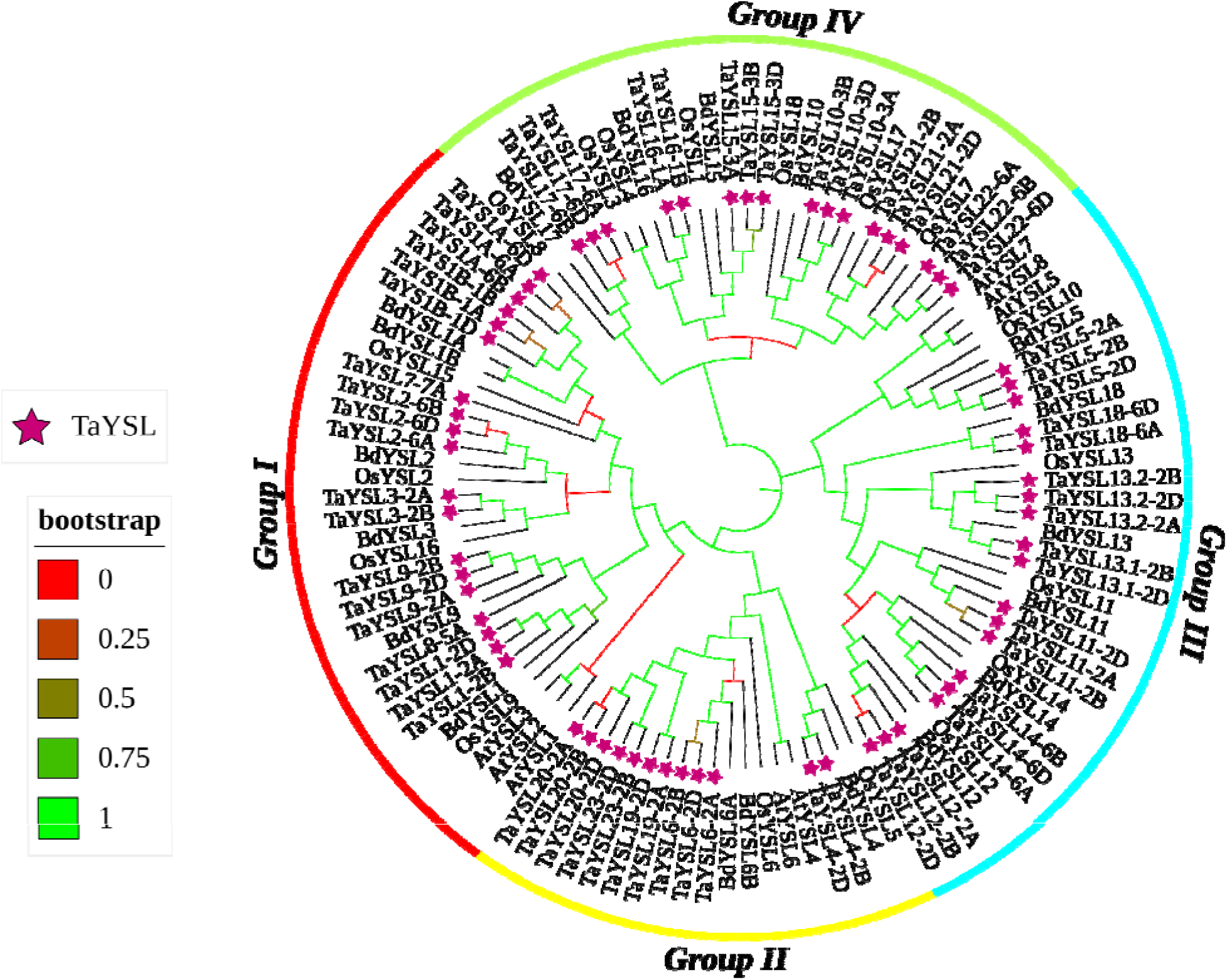
Phylogenetic tree analysis of TaYSL proteins from Arabidopsis, rice, brachypodium and wheat. Distinct color blocks correspond to various subgroups of YSL proteins. Stars represent TaYSL family members.

The exon-intron structure of TaYSL family members revealed that *TaYSL* genes consisted of 2 to 13 exons, with most members within the same subfamily exhibiting similarities in exon and intron number. However, group IV members displayed differences in exon and intron number which may contribute to functional diversification during the evolution of *TaYSL* genes. All TaYSL proteins shared core conserved domains (Pfam03169, cl14607, TIGR00728, COG1297, and cl42457), except for *TaYSL15-3B* (Group IV) and *TaYSL20-2D* (Group II), which lacked the TIGR00733 domain (Supplementary Figure S1).

The presence of abscisic acid response elements (ABREs) indicates that the majority of YSLs may function in plant respiration and in the regulatory pathway of ABA hormone due to its role in Zn and Fe transport, except *TaYS1A-6A, TaYSL5-2B* and *TaYSL14-6D*. Promoter analysis identified ABRE2 (CCACGTGG), which serves as core binding site for IRO2 in *TaYSL3-2A, TaYSL3-2B, TaYSL6-2B, TaYSL19-2A, TaYSL19-2D, TaYSL23-2B* and *TaYSL23-2D* (Supplementary Figure S2).

### Expression patterns of *TaYSL* genes in wheat

We analyzed expression data of 67 *TaYSL* genes across various wheat tissues, including root, leaf, grain, spike, and stem from datasets of cv. Chinese Spring wheat. Group I (19/67) members were highly expressed across various tissues. Among them, *TaYSL3-2B* and *TaYSL3-2A* were predominantly expressed in leaves, whereas *TaYSL9-2A*, *TaYSL9-2B*, and *TaYSL9-2D* showed high expression in roots. Most members of Group II (12/67) exhibited relatively high expression levels in roots, while *TaYSL4-2D*, *TaYSL6-2A*, *TaYSL6-2B*, and *TaYSL6-2D* displayed distinct expression patterns across different organs (Figure 2). Group III (19/67) genes were mainly expressed in grains compared with other organs, whereas Group IV (17/67) genes showed low expression in most tissues except for spikes.

**Figure 2.**
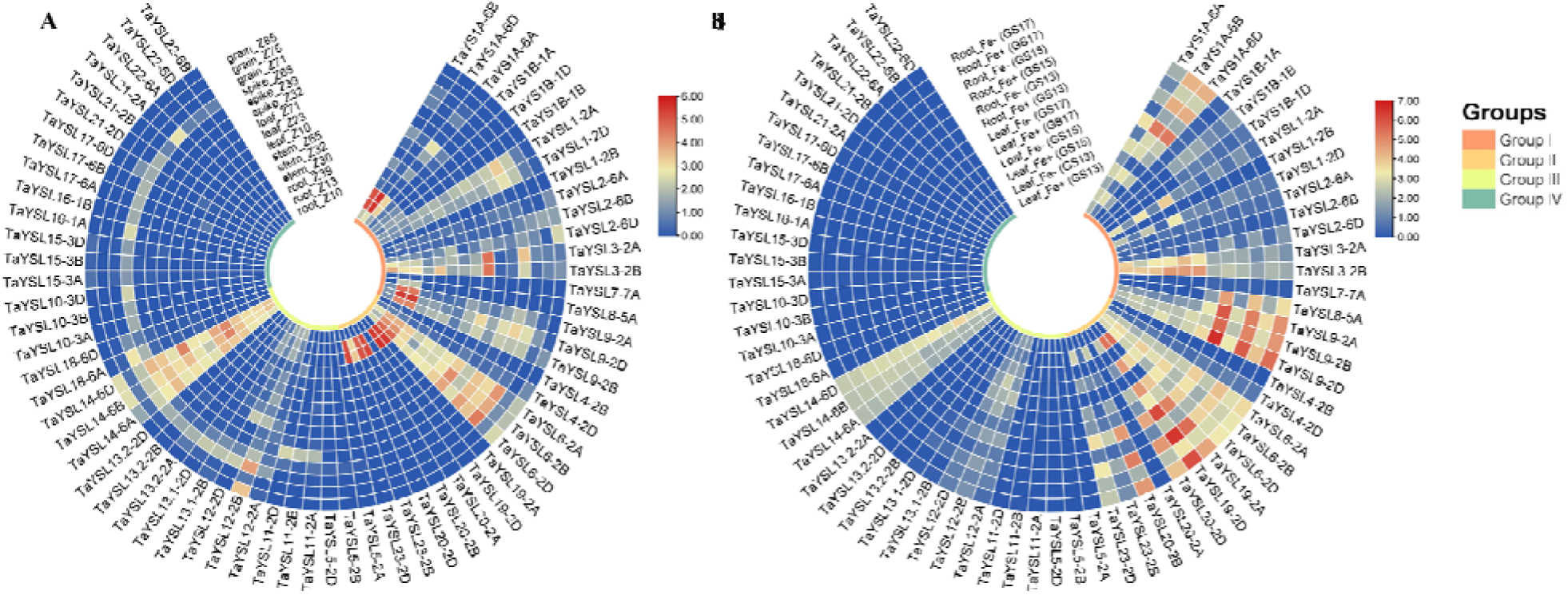
Heatmap depicts the phylogenetic grouping and expression patterns of 67 *TaYSL* genes (A) across various growth stages and tissues of Chinese Spring wheat, and (B) in roots and leaves under Fe deficiency across three developmental stages i.e., three-leaf (GS13), five-leaf (GS15), seven-leaf (GS17) stages according to the Zadok’s scale. Expression data was log2 normalized in TBtools. Blue and red represent low and high expression levels, respectively.

The expression of *TaYSL* genes under Fe deficiency was analyzed using our transcriptome resource of wheat cv. Zincol-16 (Qayyum et al., 2026). Based on RNA-seq data, 26 *TaYSL* genes showed differential expression pattern in roots and leaves across different developmental stages. Most genes belonging to Group I and Group II exhibited significant upregulation under Fe deficiency. However, the expression of *TaYSL1-2A* and *TaYSL1-2D* genes were down-regulated under Fe deficiency in leaves. Group III and Group IV exhibited no significant changes in gene expression, except *TaYSL11* and *TaYSL14* genes, indicating that genes belonging to these two groups may be tissue-specific expressed or in response to specific stress conditions. Based on RNA-seq data, five most significant DEGs (*TaYSL19, TaYSL 20, TaYSL 9, TaYSL6* and *TaYSL1*) were selected for qRT-PCR analysis, and their expression level were measured in roots and leaves at three developmental stages under optimal and Fe deficiency conditions. Expression trends from qRT-PCR validated the RNA-seq data for the selected *TaYSL* genes, with only minor discrepancies, confirming the accuracy of the transcriptome profiling (Figure 3). Our data supports the involvement of *TaYSL* genes in metal homeostasis during Fe deficiency and in mediating abiotic stress tolerance.

**Figure 3.**
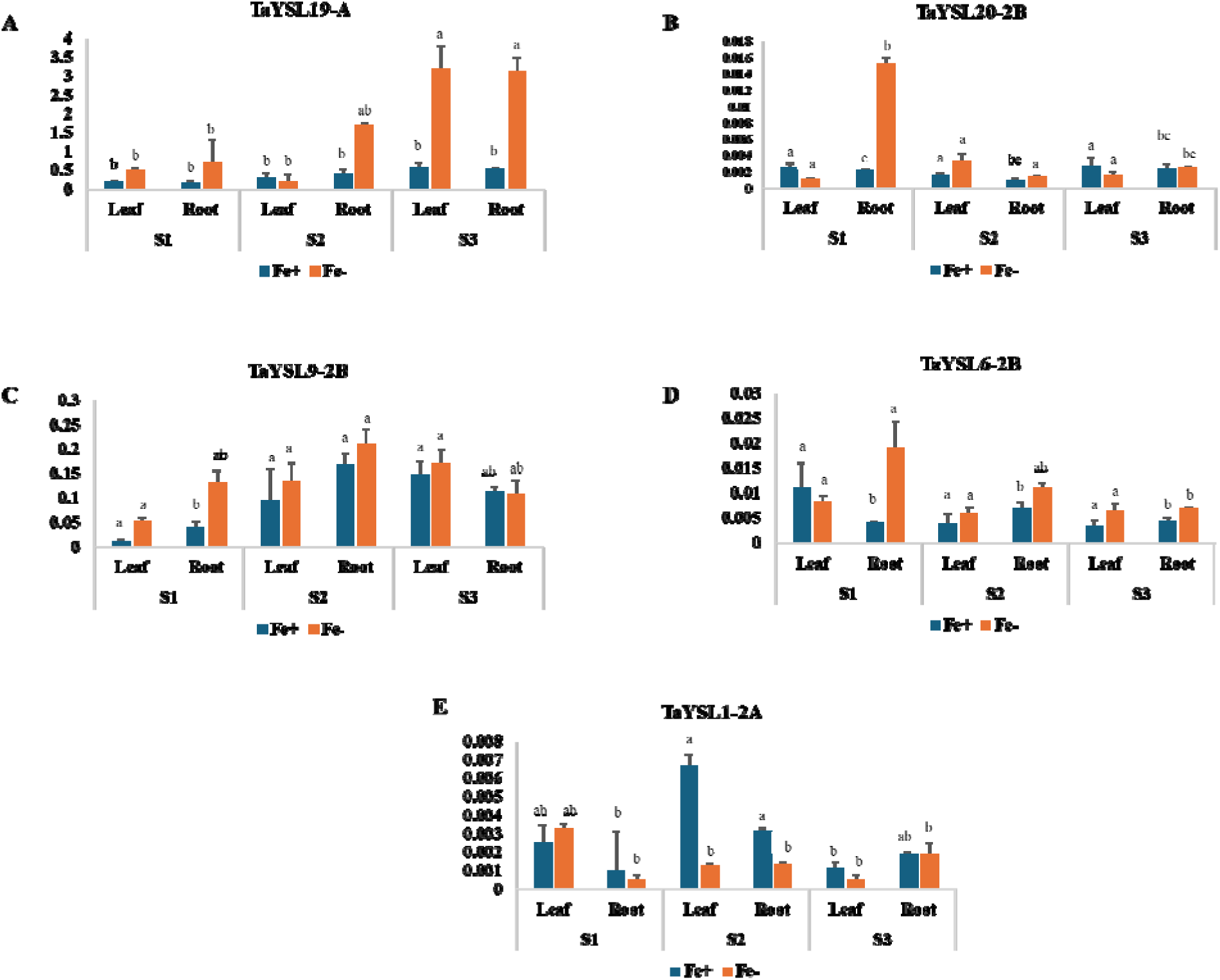
qRT-PCR analysis of (A) *TaYSL19-A, (B) TaYSL20-2B, (C) TaYSL9-2B, (D) TaYSL6- 2B, and (E) TaYSL1-2A* under Fe stress at three developmental stages: 7-day (S1), 14-day (S2) and 21-day (S3). *TaActin1* serves as the internal reference gene. Values shown are the mean ± standard error from three independent biological replicates. Statistically significant differences (p < 0.05) between means are indicated by different letters.

### Natural alleles of *TaYSLs* and their association with GFeC and GZnC in wheat

The screening of variants from 67 *TaYSL* genes identified 3346 SNPs across the genome. The A sub-genome contained the highest number of variants (n= 1762) in 23 *TaYSL* genes, followed by B (n= 1240) in 22 *TaYSL* genes, and D sub-genome (n= 344) in 16 *TaYSL* genes. Only a small fraction of SNPs were found in coding regions, with 5.8% being low-impact synonymous variants and 4.7% being moderate-impact missense variants. A high-impact stop-lost variant was identified in *TaYSL4-2B* at position 425360722 (Supplementary Figure S3). No genetic variants were detected for *TaYS1A-6D*, *TaYSL15-3D* and *TaYSL20-2D*.

Association analysis identified a total of 28 SNPs belonging to 9 YSL genes were significantly (p < 0.001) associated with GFeC in more than three out of six environments including average across environments (Table 1). Among them eight SNPs belonging to *TaYS1B-1A, TaYSL2-6B* and *TaYSL16-1A*. *TaYSL16-1A* consisted of 5 intron variants, *TaYSL2-6B* contained 2 upstream variants while *TaYS1B-1A* had one downstream gene variant, which were associated with GFeC across all six environments including their average across environments.

**Table 1.**
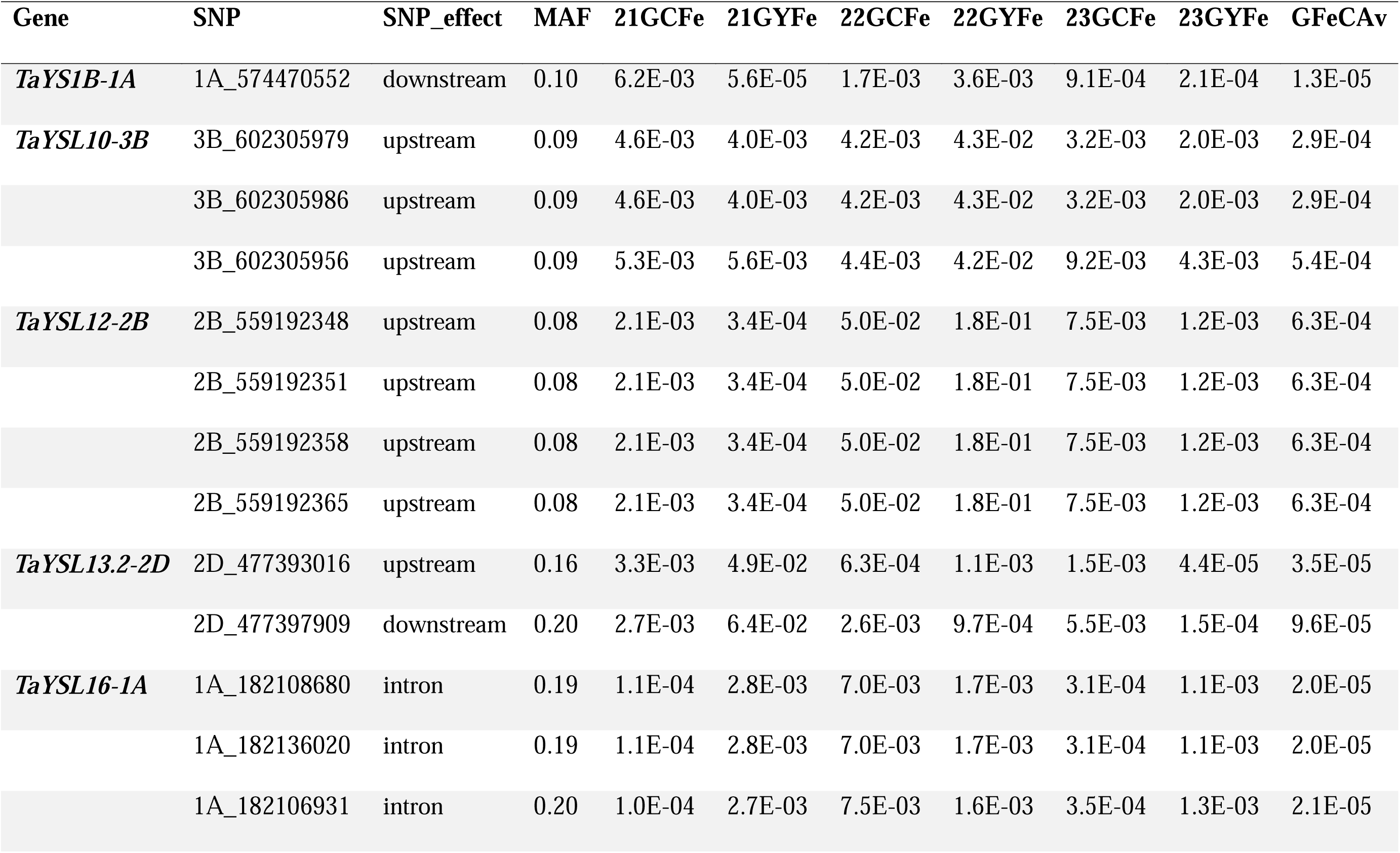
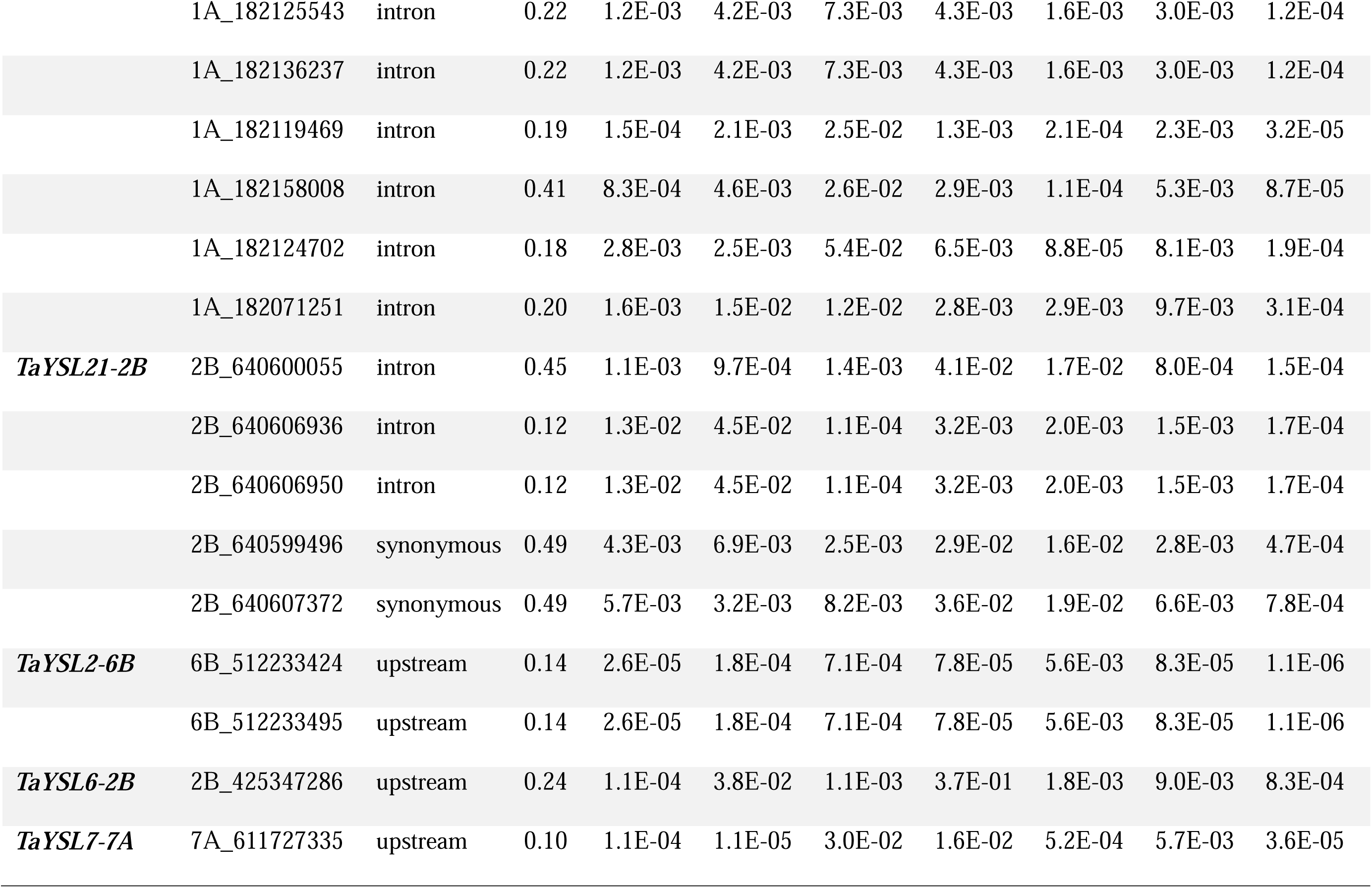
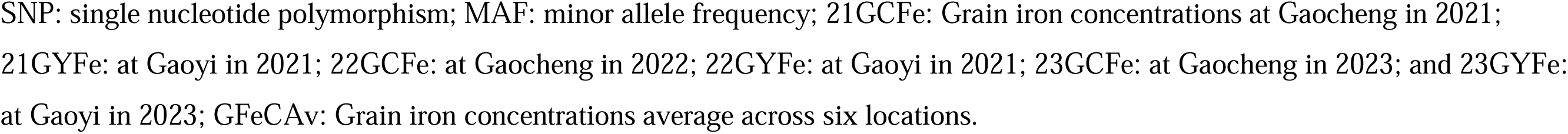
SNPs associated with GFeC in a diversity Panel of 145 cultivars grown over three years (2021-2023), at two locations in China (Gaoyi and Gaocheng).

| Gene | SNP | SNP_effect | MAF | 21GCFe | 21GYFe | 22GCFe | 22GYFe | 23GCFe | 23GYFe | GFeCAv |
| --- | --- | --- | --- | --- | --- | --- | --- | --- | --- | --- |
| **TaYS1B-1A** | 1A_574470552 | downstream | 0.10 | 6.2E-03 | 5.6E-05 | 1.7E-03 | 3.6E-03 | 9.1E-04 | 2.1E-04 | 1.3E-05 |
| **TaYSL10-3B** | 3B_602305979 | upstream | 0.09 | 4.6E-03 | 4.0E-03 | 4.2E-03 | 4.3E-02 | 3.2E-03 | 2.0E-03 | 2.9E-04 |
|  | 3B_602305986 | upstream | 0.09 | 4.6E-03 | 4.0E-03 | 4.2E-03 | 4.3E-02 | 3.2E-03 | 2.0E-03 | 2.9E-04 |
|  | 3B_602305956 | upstream | 0.09 | 5.3E-03 | 5.6E-03 | 4.4E-03 | 4.2E-02 | 9.2E-03 | 4.3E-03 | 5.4E-04 |
| **TaYSL12-2B** | 2B_559192348 | upstream | 0.08 | 2.1E-03 | 3.4E-04 | 5.0E-02 | 1.8E-01 | 7.5E-03 | 1.2E-03 | 6.3E-04 |
|  | 2B_559192351 | upstream | 0.08 | 2.1E-03 | 3.4E-04 | 5.0E-02 | 1.8E-01 | 7.5E-03 | 1.2E-03 | 6.3E-04 |
|  | 2B_559192358 | upstream | 0.08 | 2.1E-03 | 3.4E-04 | 5.0E-02 | 1.8E-01 | 7.5E-03 | 1.2E-03 | 6.3E-04 |
|  | 2B_559192365 | upstream | 0.08 | 2.1E-03 | 3.4E-04 | 5.0E-02 | 1.8E-01 | 7.5E-03 | 1.2E-03 | 6.3E-04 |
| **TaYSL13.2-2D** | 2D_477393016 | upstream | 0.16 | 3.3E-03 | 4.9E-02 | 6.3E-04 | 1.1E-03 | 1.5E-03 | 4.4E-05 | 3.5E-05 |
|  | 2D_477397909 | downstream | 0.20 | 2.7E-03 | 6.4E-02 | 2.6E-03 | 9.7E-04 | 5.5E-03 | 1.5E-04 | 9.6E-05 |
| **TaYSL16-1A** | 1A_182108680 | intron | 0.19 | 1.1E-04 | 2.8E-03 | 7.0E-03 | 1.7E-03 | 3.1E-04 | 1.1E-03 | 2.0E-05 |
|  | 1A_182136020 | intron | 0.19 | 1.1E-04 | 2.8E-03 | 7.0E-03 | 1.7E-03 | 3.1E-04 | 1.1E-03 | 2.0E-05 |
|  | 1A_182106931 | intron | 0.20 | 1.0E-04 | 2.7E-03 | 7.5E-03 | 1.6E-03 | 3.5E-04 | 1.3E-03 | 2.1E-05 |
|  | 1A_182125543 | intron | 0.22 | 1.2E-03 | 4.2E-03 | 7.3E-03 | 4.3E-03 | 1.6E-03 | 3.0E-03 | 1.2E-04 |
|  | 1A_182136237 | intron | 0.22 | 1.2E-03 | 4.2E-03 | 7.3E-03 | 4.3E-03 | 1.6E-03 | 3.0E-03 | 1.2E-04 |
|  | 1A_182119469 | intron | 0.19 | 1.5E-04 | 2.1E-03 | 2.5E-02 | 1.3E-03 | 2.1E-04 | 2.3E-03 | 3.2E-05 |
|  | 1A_182158008 | intron | 0.41 | 8.3E-04 | 4.6E-03 | 2.6E-02 | 2.9E-03 | 1.1E-04 | 5.3E-03 | 8.7E-05 |
|  | 1A_182124702 | intron | 0.18 | 2.8E-03 | 2.5E-03 | 5.4E-02 | 6.5E-03 | 8.8E-05 | 8.1E-03 | 1.9E-04 |
|  | 1A_182071251 | intron | 0.20 | 1.6E-03 | 1.5E-02 | 1.2E-02 | 2.8E-03 | 2.9E-03 | 9.7E-03 | 3.1E-04 |
| **TaYSL21-2B** | 2B_640600055 | intron | 0.45 | 1.1E-03 | 9.7E-04 | 1.4E-03 | 4.1E-02 | 1.7E-02 | 8.0E-04 | 1.5E-04 |
|  | 2B_640606936 | intron | 0.12 | 1.3E-02 | 4.5E-02 | 1.1E-04 | 3.2E-03 | 2.0E-03 | 1.5E-03 | 1.7E-04 |
|  | 2B_640606950 | intron | 0.12 | 1.3E-02 | 4.5E-02 | 1.1E-04 | 3.2E-03 | 2.0E-03 | 1.5E-03 | 1.7E-04 |
|  | 2B_640599496 | synonymous | 0.49 | 4.3E-03 | 6.9E-03 | 2.5E-03 | 2.9E-02 | 1.6E-02 | 2.8E-03 | 4.7E-04 |
|  | 2B_640607372 | synonymous | 0.49 | 5.7E-03 | 3.2E-03 | 8.2E-03 | 3.6E-02 | 1.9E-02 | 6.6E-03 | 7.8E-04 |
| **TaYSL2-6B** | 6B_512233424 | upstream | 0.14 | 2.6E-05 | 1.8E-04 | 7.1E-04 | 7.8E-05 | 5.6E-03 | 8.3E-05 | 1.1E-06 |
|  | 6B_512233495 | upstream | 0.14 | 2.6E-05 | 1.8E-04 | 7.1E-04 | 7.8E-05 | 5.6E-03 | 8.3E-05 | 1.1E-06 |
| **TaYSL6-2B** | 2B_425347286 | upstream | 0.24 | 1.1E-04 | 3.8E-02 | 1.1E-03 | 3.7E-01 | 1.8E-03 | 9.0E-03 | 8.3E-04 |
| **TaYSL7-7A** | 7A_611727335 | upstream | 0.10 | 1.1E-04 | 1.1E-05 | 3.0E-02 | 1.6E-02 | 5.2E-04 | 5.7E-03 | 3.6E-05 |

SNP: single nucleotide polymorphism; MAF: minor allele frequency; 21GCFE: Grain iron concentrations at Gaocheng in 2021; 21GYFe: at Gaoyi in 2021; 22GCFE: at Gaocheng in 2022; 22GYFe: at Gaoyi in 2021; 23GCFE: at Gaocheng in 2023; and 23GYFe: at Gaoyi in 2023; GFeCAv: Grain iron concentrations average across six locations.

53 SNPs belonging to *TaYSL4-2B*, *TaYSL6-2B*, *TaYSL14-6A*, *TaYSL16-1A* and *TaYSL18-6A* were significantly associated with GZnC in more than three environments including average. *TaYSL14-6A*, *TaYSL6-2B* and *TaYSL18-6A* had nineteen, one and thirteen SNPs associated with GZnC in more than five environments including average across environments (Table S4). Among all the genes analyzed, intronic variants of *TaYSL16-1A* were significantly associated with both GFeC and GZnC across different environments.

RTM-GWAS divided 3346 SNPs into 241 haplotypes, grouping SNPs with the same linkage disequilibrium (LD) block into a haplotype. A total of four haplotypes were identified, including *TaYSL2-6B*, *TaYSL6-2B*, *TaYSL7-7A* and *TaYSL16-1A* which showed significant association with GFeC and *TaYSL14-6A* with GZnC. Two SNPs on chr6B formed two haplotypes while a significant cluster of 14 SNPs was found on chr2B forming six haplotypes. Additionally, 6 SNPs on chr7A formed seven haplotypes. Haplotypes present in fewer than 10 varieties were discarded while the phenotypic variation among different haplotypes ranged from 39-56 mg kg^−1^. The haplotypes of *TaYSL6-2B*, *TaYSL16-1A* and *TaYSL14-6A* are shown in Figure 4-6, while the haplotypes of *TaYSL2-6B* and *TaYSL7-7A* are shown in Supplementary Figures S4 and S6.

**Figure 4.**
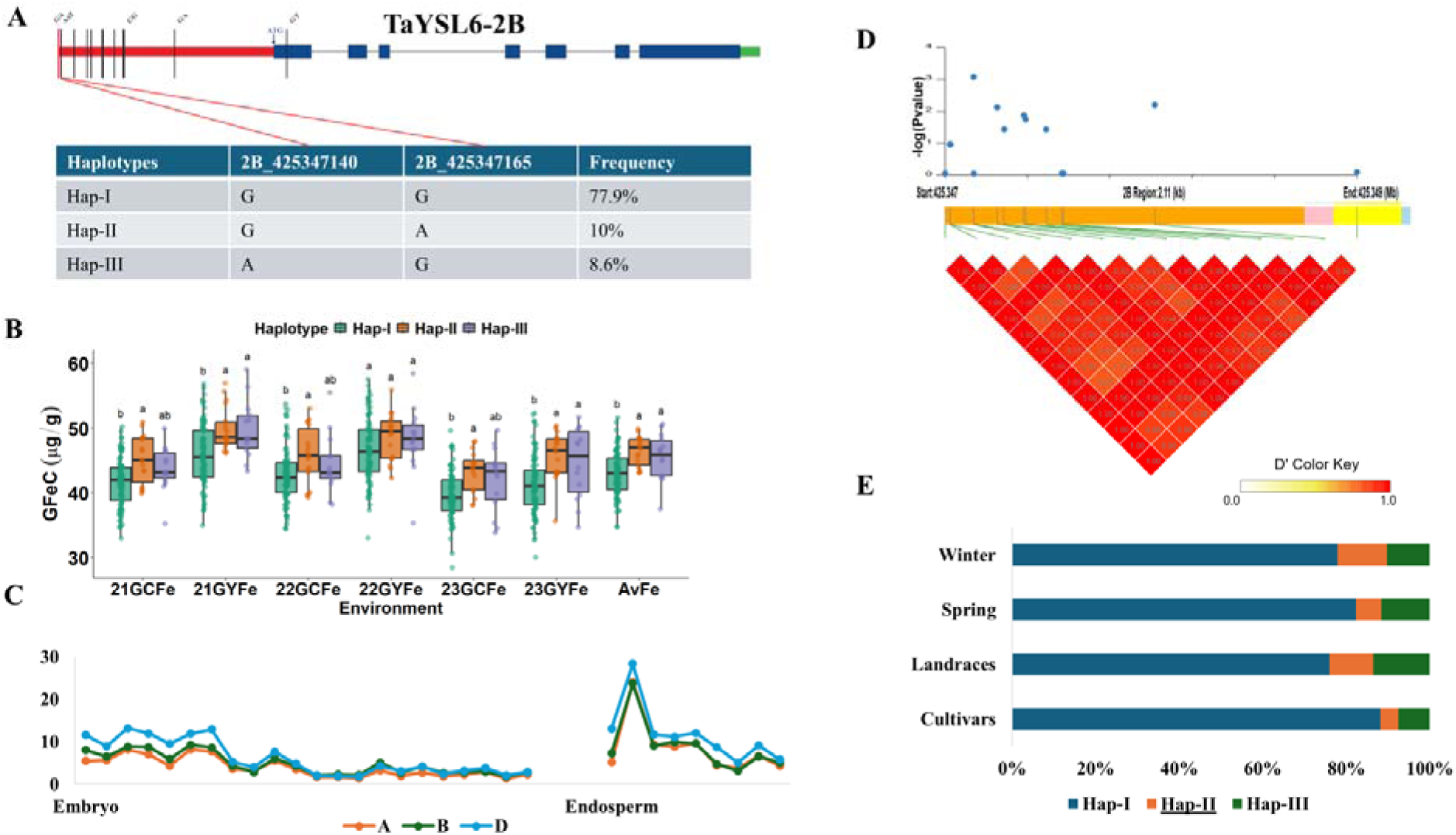
Haplotype analysis of *TaYSL6-2B* and its association with grain iron concentration. (A) Gene structure displays polymorphic sites and identified haplotypes (Hap-I, Hap-II, and Hap-III), along with their frequencies in the population. (B) Comparison of grain iron concentration among the three haplotypes across six different environments including average. (C) Expression pattern of each homeolog of *TaYSL6* in embryo and endosperm tissues. (D) LD heatmap illustrating LD patterns within the gene region. (E) Frequency of the favorable haplotype (underlined) in global wheat collections categorized into landraces, cultivars, winter and spring wheat.

**Figure 5.**
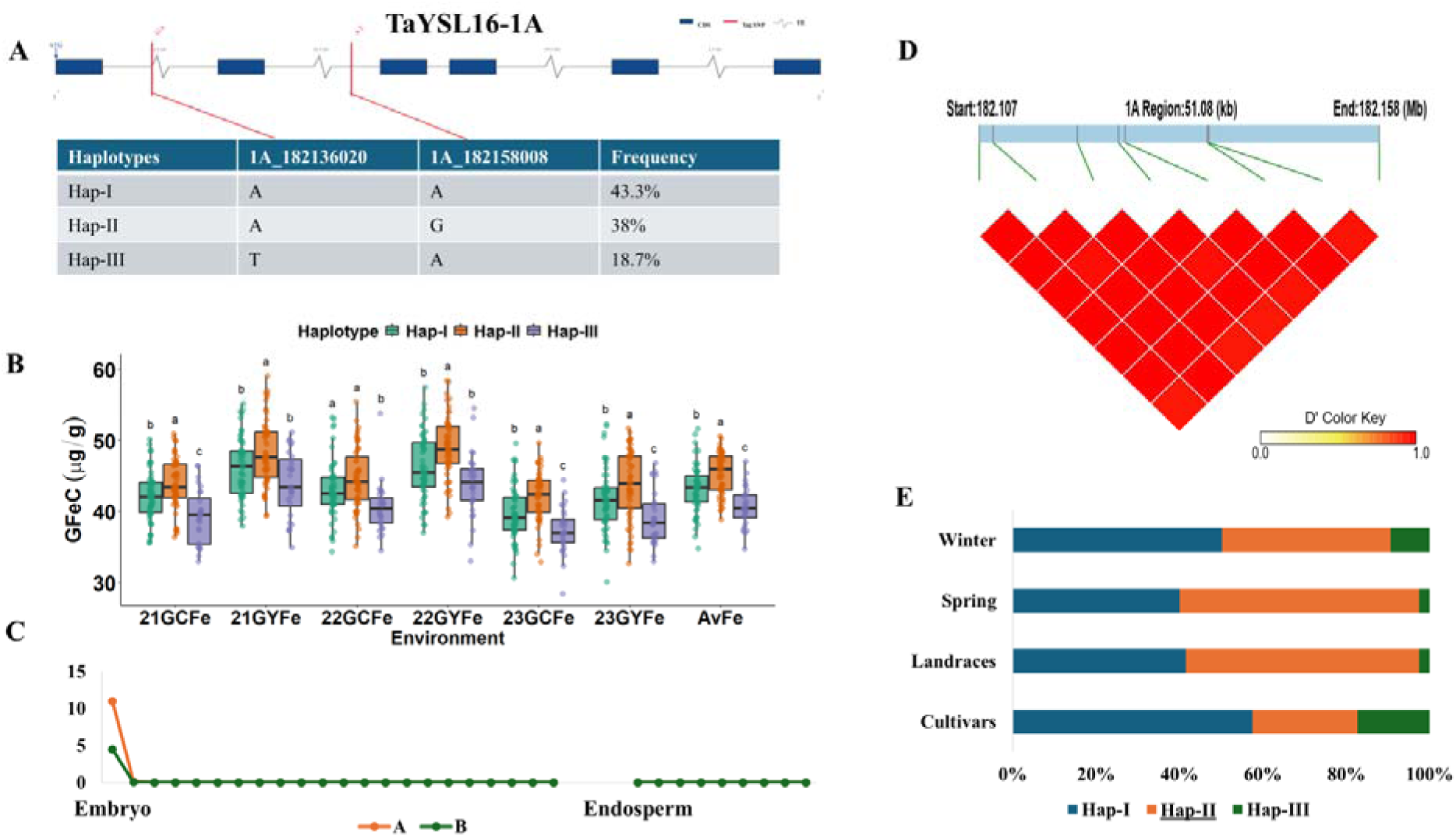
Haplotype analysis of *TaYSL16-1A* and its association with grain iron concentration. (A) Gene structure displays polymorphic sites and identified haplotypes (Hap-I, Hap-II, and Hap-III), along with their frequencies in the population. (B) Comparison of grain iron concentration among the three haplotypes across six different environments including average. (C) Expression pattern of each homeolog of *TaYSL16* in embryo and endosperm tissues. (D) LD heatmap illustrating LD patterns within the gene region. (E) Frequency of the favorable haplotypes (underlined) in global wheat collections categorized into landraces, cultivars, winter and spring wheat.

**Figure 6.**
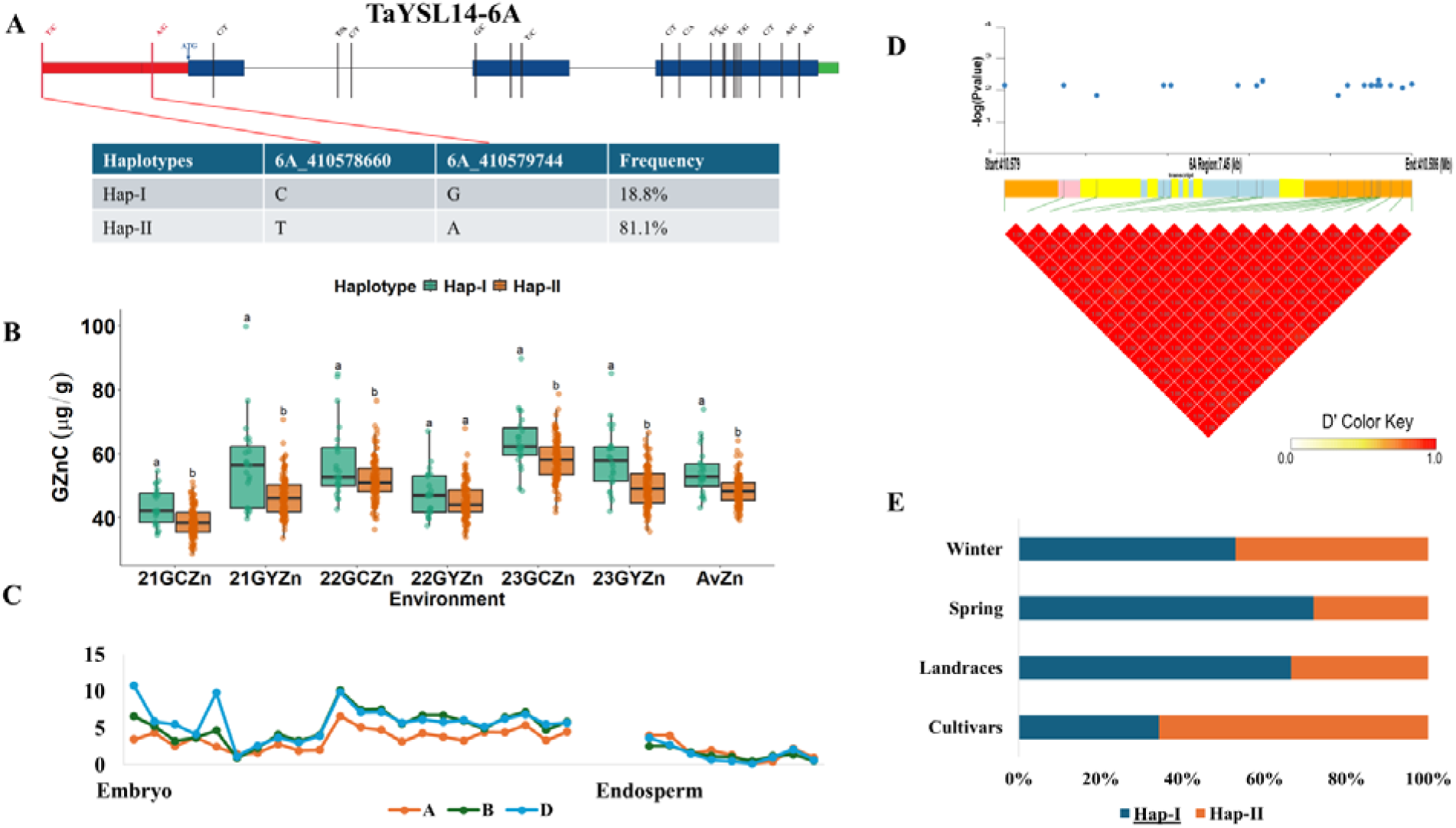
Haplotype analysis of *TaYSL14-6A* and its association with grain zinc concentration. (A) Gene structure displays polymorphic sites and identified haplotypes (Hap-I and Hap-II), along with their frequencies in the population. (B) Comparison of grain zinc concentration among the two haplotypes across six different environments including average. (C) Expression pattern of each homeolog of *TaYSL14* in embryo and endosperm tissues. (D) LD heatmap illustrating LD patterns within the gene region. (E) Frequency of the favorable haplotype (underlined) in global wheat collections categorized into landraces, cultivars, winter and spring wheat.

### Grain-specific expression and allele frequency distribution in global wheat germplasm

Based on the spatiotemporal transcriptomic landscape of wheat embryo and endosperm development (Guo et al., 2025), the expression patterns of candidate genes significantly associated with GFeC and GZnC were examined in both tissues. Among all the genes analyzed *TaYSL2-6B, TaYSL6-2B* and *TaYSL14-6* had high expression in endosperm suggesting their distinct roles during seed development. *TaYSL7-7A* and *TaYSL16-1A* had embryo-specific expression only. *TaYSL16-1A* had no expression in endosperm and was only expressed in embryo at initial stages while its favorable haplotype was present in 40.7% of land races and 9.2% of cultivars. The favorable haplotypes of *TaYSL2-6B* and *TaYSL7-7A* were in more than 80% of landraces and cultivars while the favorable haplotype of *TaYSL6-2B* was limited and present in 43.8% of landraces and 6.6% of cultivars.

### Distribution of favorable haplotypes in global wheat collections

We further calculated the frequency of haplotype combinations of three favorable genes across a global wheat collection of ∼3000 accessions (Figure 7). We obtained eight different combinations based on the presence of positive and neutral alleles of three YSL genes. The favorable haplotype combination at three genes i.e. TaYSL6^P^, TAYSL16^P^, TaYSL14^P^, was relatively rare representing 2.7% (65 accession) of the total accessions. The favorable haplotype was most frequent in Chinese Landraces (5.1%), followed by European Winter Wheat (3.7%) while the lowest frequency was observed in Watkins Landraces (1.6%) and Chinese cultivars (1.7%).

**Figure 7.**
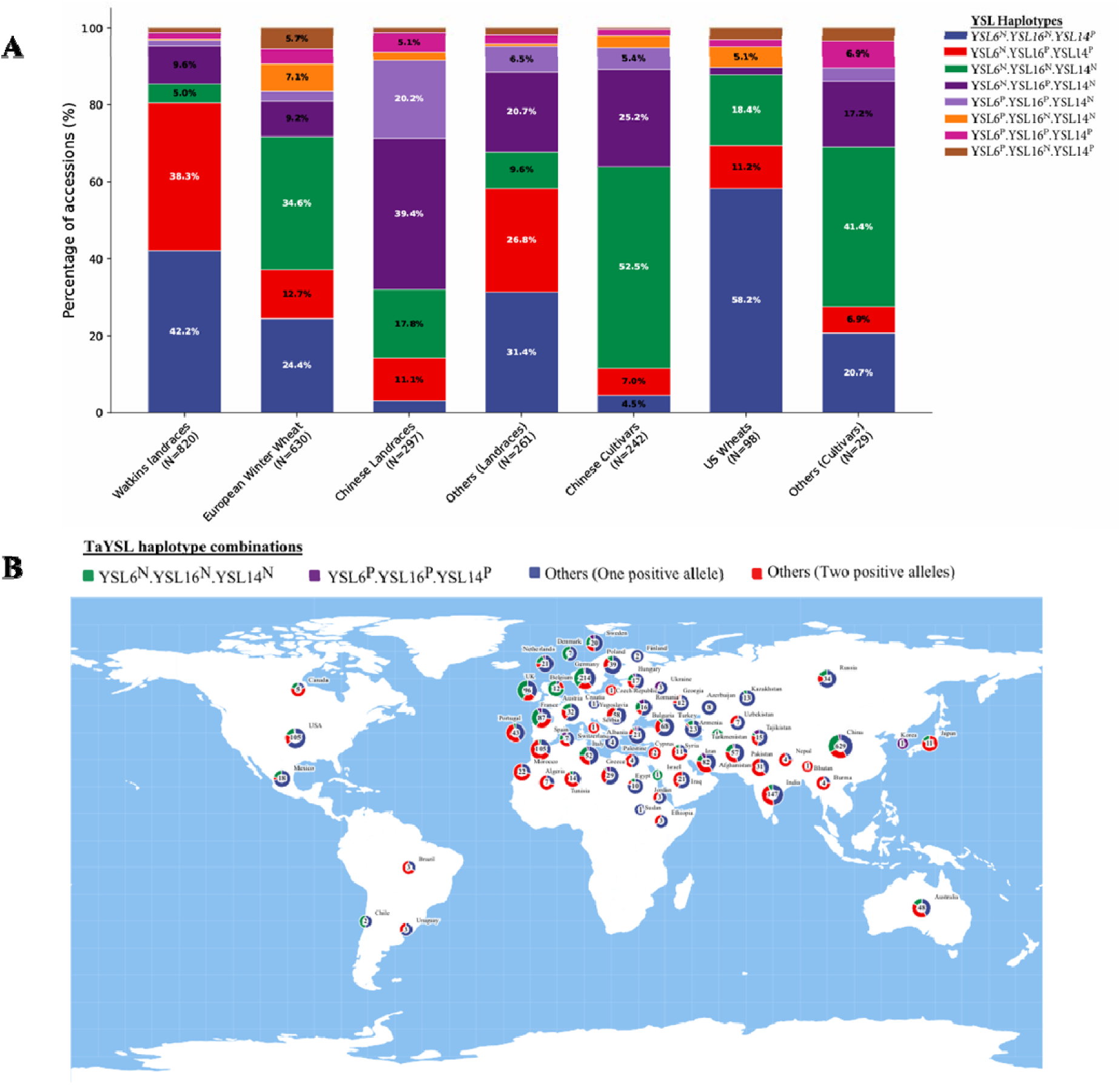
(A) Stacked bar plot illustrates percentage of different *YSL* haplotypes across various wheat groups including Watkins landraces, European Winter Wheat, Chinese, US and other landraces and cultivars showing diversity within each. (B) The world map represents geographic distribution of two *TaYSL* haplotype combinations. P represents positive allele while N denote Neutral Allele.

## Discussion

In this study, we systematically identified 26 *TaYSL* genes, representing 67 homoeologs in wheat. This refinement of the *TaYSL* gene family, based on the updated IWGSC RefSeq v1.1 assembly, improved annotation accuracy by removing redundant and incomplete sequences, thereby ensuring more reliable downstream analyses. The A, B, and D sub-genomes contained 23, 22, and 22 members respectively, which is greater than that of other model species. This expansion may be due to the large genome size and higher ploidy level of wheat, as well as the presence of duplicate genes that likely originated through its two-step evolutionary history (Walkowiak et al., 2020; Glover et al., 2015).

The domain architecture and motif organization of TaYSL proteins showed high conservation among members of the same phylogenetic clade, aligning with findings from earlier studies (Song et al., 2024). Genetic structural diversification significantly contributes to the evolution of multi-gene families (Muthamilarasan et al., 2014; Han et al., 2016). The *TaYSL* genes exhibited variation in exon-intron structure, potentially arising from genomic fragment integration and subsequent rearrangement (Xu et al., 2012). The phylogenetic tree results showed that these 67 TaYSL proteins divided YSLs into four groups consistent with the earlier reports (Koike et al., 2004; Gross et al., 2003; Yordem et al., 2011). Despite being non-hyperaccumulators, rice and wheat exhibit evolutionary conservation in their heavy metal uptake and translocation mechanisms (Liu et al., 2025). In rice, Fe deficiency regulates *YSL* gene expression through multiple transcription factors. Among these, IRO2 acts as a key regulator by directly interacting with iron-responsive elements (IREs) in promoter regions to activate transcription (Saleem et al., 2025). In wheat, we identified an IRO2 binding site in group I and group II members highlighting their role in Fe homeostasis under Fe deficiency.

*YSL* genes showed tissue-specific expression profiles, showing variable transcript abundance in roots, leaves, spikes, and grains. The tissue-specific expression profiles of *YSL* genes correlate with the multifunctional roles of YSL transporters in Fe acquisition, transport and distribution processes (Saleem et al., 2024). Expression analysis indicated that *TaYSL* genes are generally upregulated under abiotic stress. In wheat, group I and II exhibited differential expressions in response to Fe deficiency in both roots and leaves. These findings demonstrate the metal selectivity of these transporters, representing a key limitation for enhancing grain Fe and Zn accumulation (Sharma et al., 2020). *OsYSL9* showed substantial upregulation at higher Fe concentrations, underscoring its sensitivity to environmental Fe levels (Saleem et al., 2004). In wheat, its orthologue *TaYSL1* demonstrated a similar expression pattern, suggesting a negative regulatory role in Fe homeostasis. Moreover, its knockout mutants showed significant increase in both Fe and Zn content which further confirms it as a repressor of nutrient uptake. Disrupting these genes through genome editing eliminates their natural inhibitory effect, leading to increased micronutrient accumulation.

According to studies, the first subgroup participates in metal transport, while the second subgroup regulates excess accumulation and detoxification, together forming a relatively complete internal and external cycle of Fe homeostasis (Song et al., 2024). Despite having a wider range of metal substrate specificity, *YSL* genes are primarily responsible for moving Fe-chelates across the plasma membrane (Kumar et al., 2025). Hence, a greater number of genes identified in our analysis showed significantly stronger associations with Fe compared to Zn. Genes with significant SNPs in coding or regulatory regions, such as *TaYSL2-6B, TaYSL6-2B, TaYSL12-2B* and *TaYSL14-6A* also play a key role in iron stress, suggesting their involvement at the transcriptional level. Similarly, the expression of these genes peaked in spikes and grain, indicating a potential role in remobilization of nutrients into grain.

Previously, *TaYSL12-6B* was identified as a strong candidate for grain iron biofortification (Qayyum et al., 2026). In this study, upstream variants of same gene were identified as significantly associated across four different environments. All SNPs identified were in perfect linkage (r^2^=1) with each other and therefore carried identical genetic information. To capture greater genetic variation and LD information, haplotype analysis was performed. Our analysis showed that haplotypes of *TaYSL2-6B, TaYSL6-2B, TaYSL7-7A* and *TaYSL16-1A* were significantly associated with GFeC, while *TaYSL14-6A* was associated with GZnC.

*TaYSL6-2B* (Hap-II) was associated with 29.7% increase in GFeC, was present in 43.8% of landraces and 6.6% of cultivars. *TaYSL16-1A* (Hap-II) showed 10.6% increase in GFeC while 15.1% increase in GZnC content and was present in 40.7% of landraces and 9.2% of cultivars. *TaYSL14-6A (*Hap-I) associated with 12.4% increase in GZnC, was present in 55.9% landraces and 25.2% cultivars. Overall, the favorable haplotypes of each gene were more common in landraces than in modern cultivars implying that beneficial alleles may not have undergone strong positive selection during breeding. Higher expression of *TaYSL6-2B*, *TaYSL14-6A* in the endosperm indicates their potential in enhancing micronutrient accumulation in the edible part of the grain. Similarly, among the eight different haplotype combinations of *TaYSL6*, *TAYSL16*, *TaYSL14*, the favorable stacked haplotype (TaYSL6^P^, TAYSL16^P^, TaYSL14^P^) was rare across global wheat collection. Its high frequency in Chinese landraces and European winter wheat while its low frequency in Watkins landraces and Chinese cultivars suggests that traditional germplasm retains untapped genetic variation. As major breeding goal for enhancing crop nutrition is raising GZnC and GFeC. Pyramiding these haplotypes through marker-assisted selection represents a promising strategy for improving grain iron and zinc concentration in wheat.

## Conclusion

This paper systematically characterizes wheat *YSL* gene family, including genome-wide identification, structural analysis, expression profiling and haplotype analysis. Twenty-six *TaYSL* genes were identified and classified into four conserved subfamilies. Expression analysis confirmed the roles of *YSL* genes in iron regulation, with varied expression patterns under iron-deficient conditions. Combining SNP and haplotype-based GWAS across six environments, we identified haplotypes of *TaYSL2-6B, TaYSL6-2B, TaYSL7-7A, TaYSL16-1A* and *TaYSL14-6A* as key regulators of grain Fe and Zn levels. The most favorable haplotypes of *TaYSL6-2B, TaYSL16-1A* and *TaYSL14-6A* as well as their combined haplotype were rare in modern cultivars indicating their potential in breeding. Overall, these findings suggest that pyramiding elite haplotypes of YSL will be effective in wheat breeding programs to improve GFeC and GZnC for developing nutritionally enhanced cultivars through marker-assisted selection.

## List of abbreviations

YSL: Yellow Stripe-Like
TaYSL: Triticum aestivum
SNP: Single Nucleotide Polymorphism
GFeC: Grain Iron Content
GZnC: Grain Zinc Content
Fe: Iron
Zn: Zinc
IWGSC: International Wheat Genome Sequencing Consortium
TPM: Transcripts Per Kilobase Million
qRT-PCR: Quantitative Real-Time Polymerase Chain Reaction
GLM-GWAS: General Linear Model GWAS
RTM-GWAS: Restricted Two-Stage Multi-locus GWAS
LD: Linkage Disequilibrium
MAF: Minor Allele Frequency
VCF: Variant Call Format
MEME: Multiple Expectation Maximization for Motif Elicitation
GSDS: Gene Structure Display Server

## Declarations

### Ethics approval and consent to participate

Not applicable

### Consent for publication

Not applicable

### Authors contributions

KA conducted analysis and wrote the draft of the manuscript, KA, HQ, SN, MS, YH and YD performed data analysis, ZH, AW reviewed and edited the writing of the manuscript. All authors read and approved the final manuscript.

## Acknowledgements

We appreciate the scientist’s contributions to the Wheat Union database, which contains a substantial amount of whole genome resequencing data.

## Supplementary information

Table S1. List of the cultivars used in this study

Table S2. Gene-specific primers for qRT-PCR

Table S3. TaYSL family members in wheat

Table S4. Association analysis of SNPs in the *TaYSL* genes in wheat cultivars panel with three year’s (2021-2023) Fe and Zn grain data of WGRS cultivars grown at Gaoyi (GY), and Gaocheng (GC) locations (China). Yellow indicating p < 0.05 and green indicating p < 0.01.

