## Supplementary Figure for "Haplotypes variations of yellow stripe like (*TaYSL*) genes are associated with grain iron and zinc contents in wheat (*Triticum aestivum L.*)": Supplementary File 1.pptx

### Slide 1
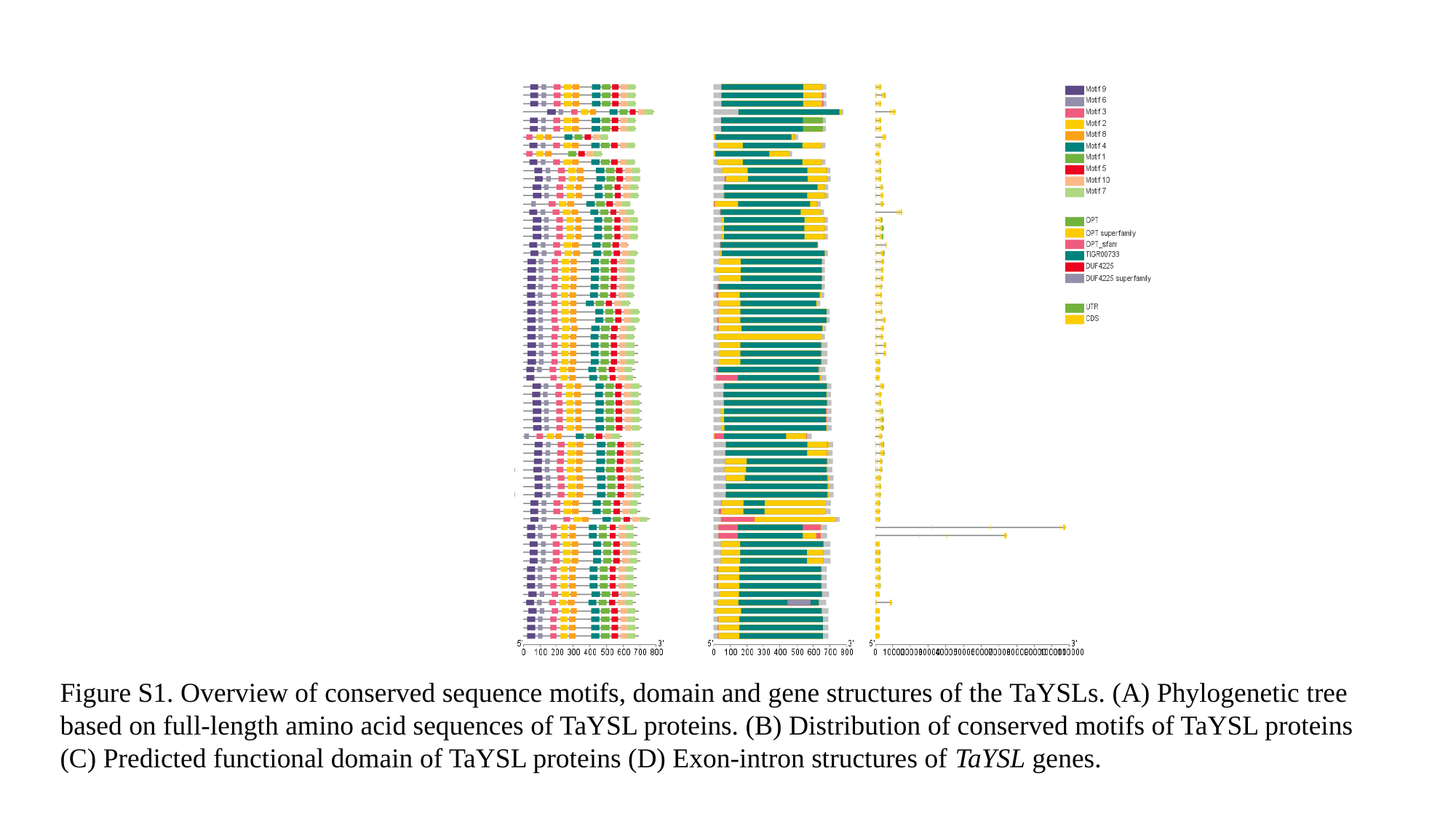

Figure S1. Overview of conserved sequence motifs, domain and gene structures of the TaYSLs. (A) Phylogenetic tree based on full-length amino acid sequences of TaYSL proteins. (B) Distribution of conserved motifs of TaYSL proteins (C) Predicted functional domain of TaYSL proteins (D) Exon-intron structures of TaYSL genes.

### Slide 2
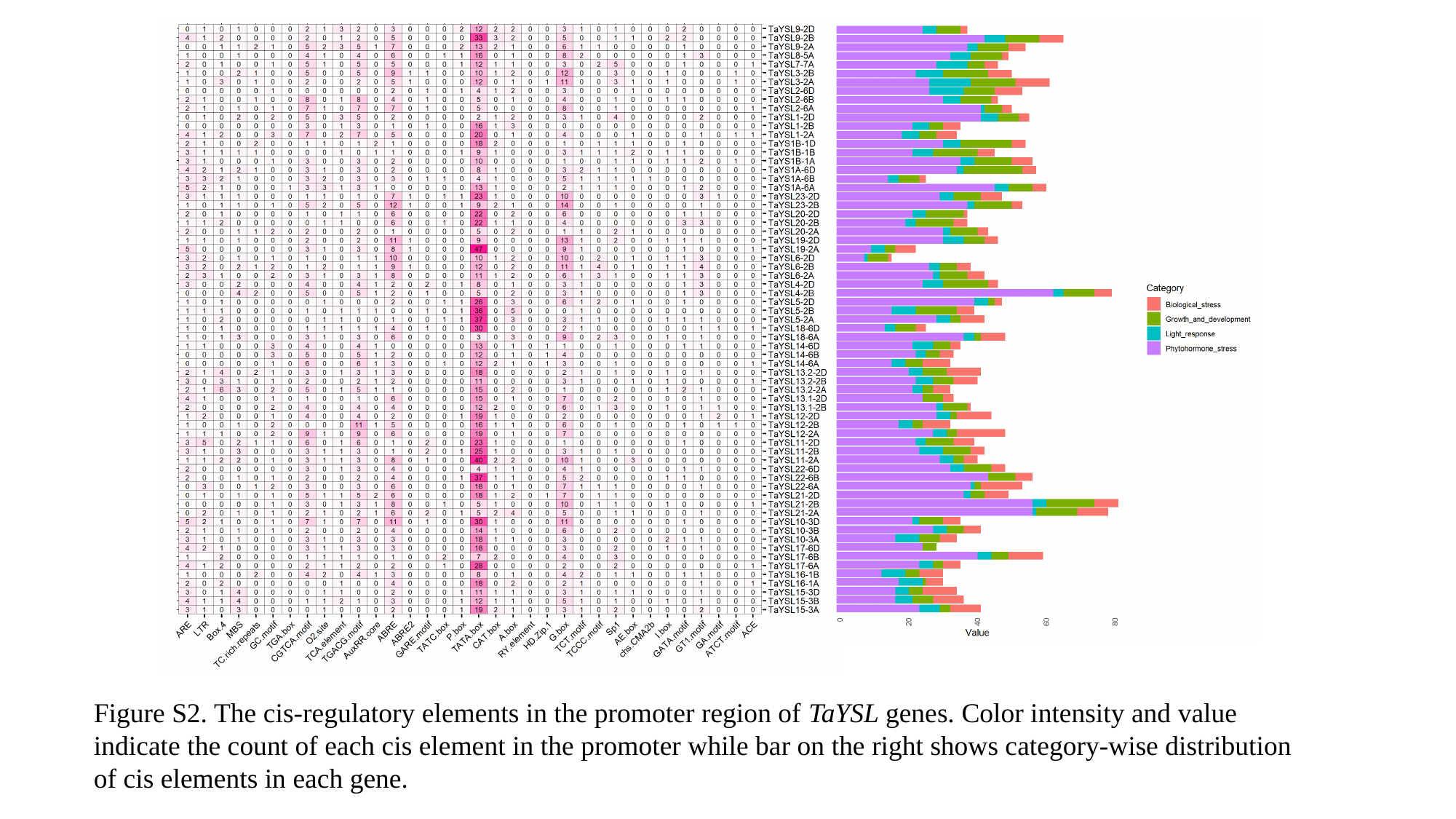

Figure S2. The cis-regulatory elements in the promoter region of TaYSL genes. Color intensity and value indicate the count of each cis element in the promoter while bar on the right shows category-wise distribution of cis elements in each gene.

### Slide 3
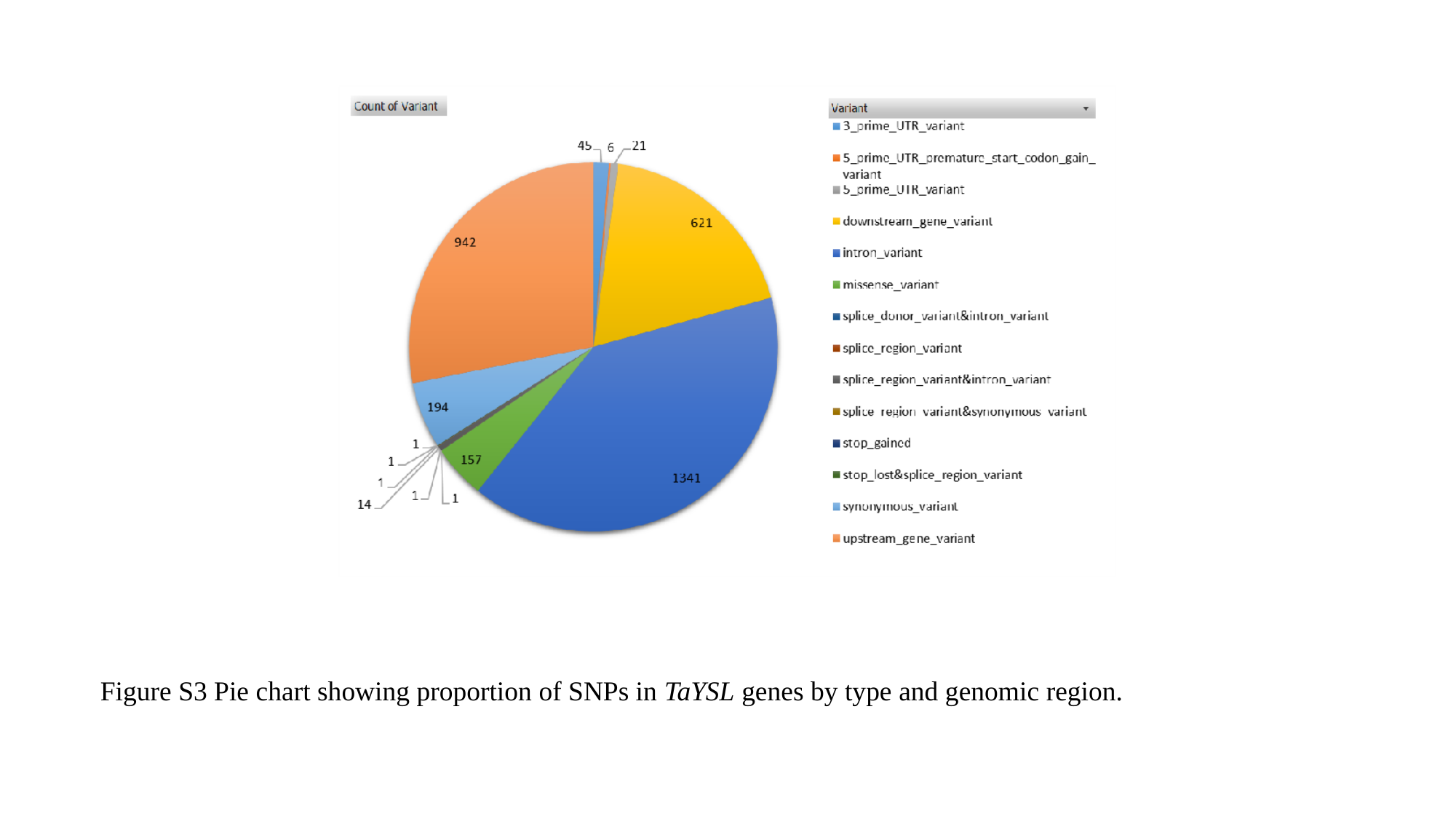

Figure S3 Pie chart showing proportion of SNPs in TaYSL genes by type and genomic region.

### Slide 4
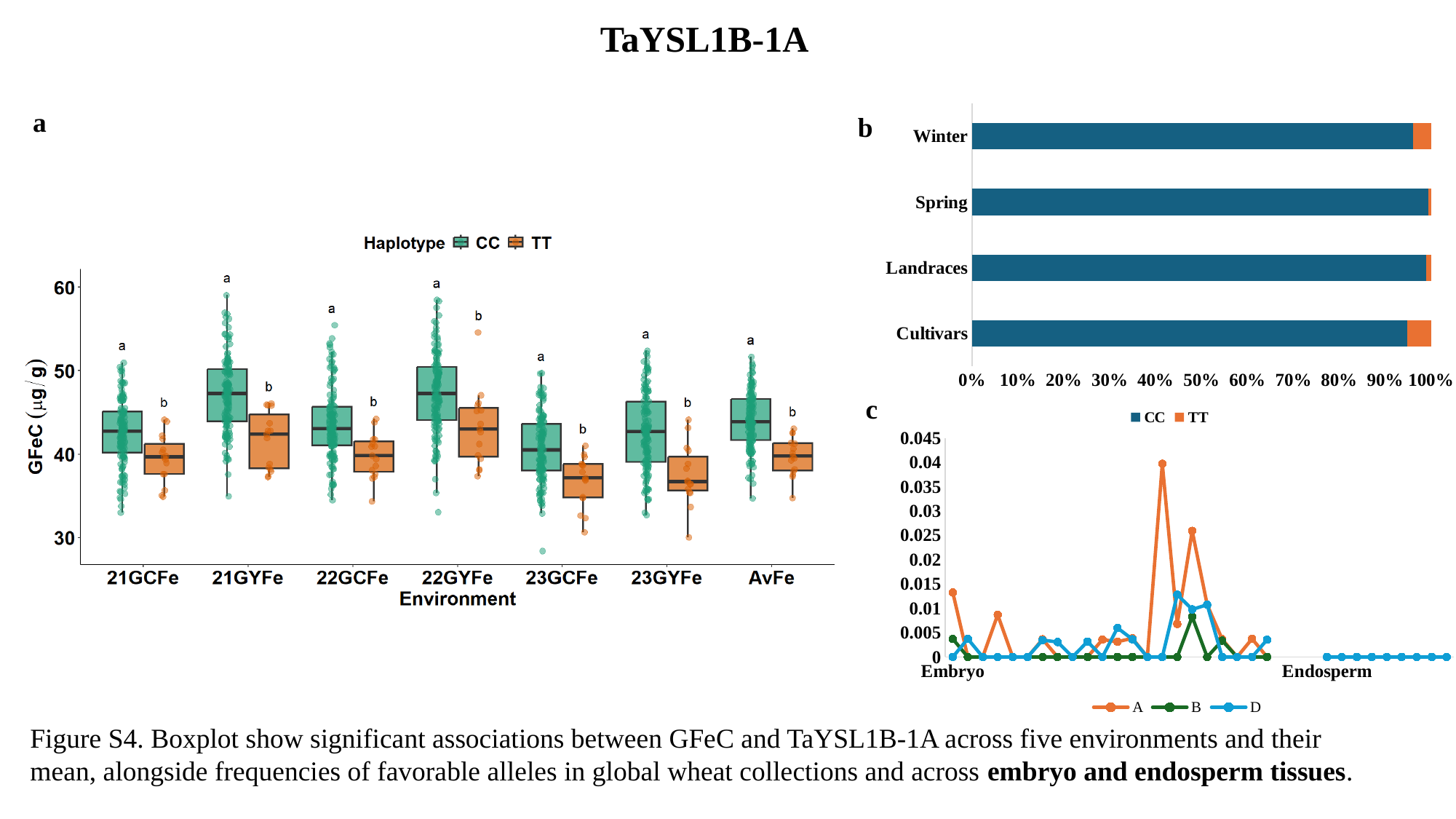

TaYSL1B-1A
#### Chart
| Category | CC | TT |
|---|---|---|
| Cultivars | 845.0 | 46.0 |
| Landraces | 746.0 | 8.0 |
| Spring | 181.0 | 1.0 |
| Winter | 1305.0 | 52.0 |a
b
c
#### Chart
| Category | A | B | D |
|---|---|---|---|
| Embryo | 0.0132333 | 0.0037 | 0.0 |
| | 0.0 | 0.0 | 0.00373333 |
| | 0.0 | 0.0 | 0.0 |
| | 0.00866667 | 0.0 | 0.0 |
| | 0.0 | 0.0 | 0.0 |
| | 0.0 | 0.0 | 0.0 |
| | 0.00363333 | 0.0 | 0.00346667 |
| | 0.0 | 0.0 | 0.00306667 |
| | 0.0 | 0.0 | 0.0 |
| | 0.0 | 0.0 | 0.00316667 |
| | 0.00356667 | 0.0 | 0.0 |
| | 0.00313333 | 0.0 | 0.00596667 |
| | 0.00383333 | 0.0 | 0.00363333 |
| | 0.0 | 0.0 | 0.0 |
| | 0.0397333 | 0.0 | 0.0 |
| | 0.00676667 | 0.0 | 0.0128 |
| | 0.0259333 | 0.00826667 | 0.00976667 |
| | 0.0107333 | 0.0 | 0.0107333 |
| | 0.00366667 | 0.00336667 | 0.0 |
| | 0.0 | 0.0 | 0.0 |
| | 0.00373333 | 0.0 | 0.0 |
| | 0.0 | 0.0 | 0.00353333 |
| | None | None | None |
| | None | None | None |
| | None | None | None |
| Endosperm | 0.0 | 0.0 | 0.0 |Figure S4. Boxplot show significant associations between GFeC and TaYSL1B-1A across five environments and their mean, alongside frequencies of favorable alleles in global wheat collections and across embryo and endosperm tissues.

### Slide 5
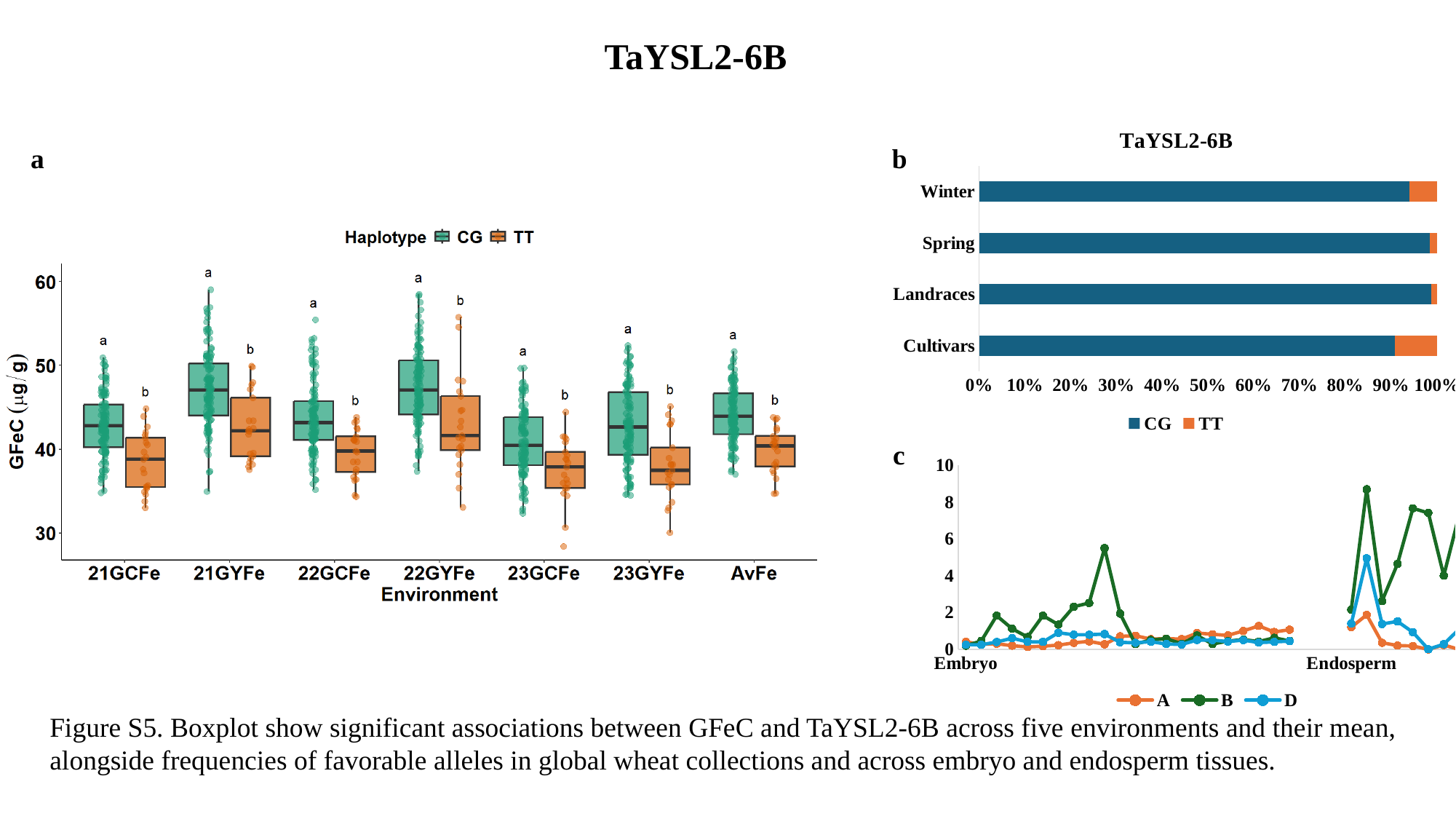

TaYSL2-6B
#### Chart: TaYSL2-6B
| Category | CG | TT |
|---|---|---|
| Cultivars | 630.0 | 64.0 |
| Landraces | 648.0 | 8.0 |
| Spring | 127.0 | 2.0 |
| Winter | 1066.0 | 67.0 |a
b
c
#### Chart
| Category | A | B | D |
|---|---|---|---|
| Embryo | 0.409667 | 0.201133 | 0.252033 |
| | 0.271833 | 0.4566 | 0.252067 |
| | 0.305567 | 1.83073 | 0.4 |
| | 0.2029 | 1.12323 | 0.606567 |
| | 0.1307 | 0.6717 | 0.409833 |
| | 0.170933 | 1.83453 | 0.405667 |
| | 0.2289 | 1.34417 | 0.902367 |
| | 0.354367 | 2.3164 | 0.796267 |
| | 0.4374 | 2.52117 | 0.797067 |
| | 0.275967 | 5.5009 | 0.829633 |
| | 0.708267 | 1.94467 | 0.384567 |
| | 0.740633 | 0.291433 | 0.355867 |
| | 0.556167 | 0.511133 | 0.418133 |
| | 0.58575 | 0.5835 | 0.2906 |
| | 0.557467 | 0.301433 | 0.2681 |
| | 0.8876 | 0.753533 | 0.508533 |
| | 0.817933 | 0.290267 | 0.5037 |
| | 0.760467 | 0.448567 | 0.423067 |
| | 1.00333 | 0.532733 | 0.502467 |
| | 1.27287 | 0.423667 | 0.366967 |
| | 0.9456 | 0.626567 | 0.4043 |
| | 1.0686 | 0.461067 | 0.450067 |
| | None | None | None |
| | None | None | None |
| | None | None | None |
| Endosperm | 1.2049 | 2.1633 | 1.3965 |Figure S5. Boxplot show significant associations between GFeC and TaYSL2-6B across five environments and their mean, alongside frequencies of favorable alleles in global wheat collections and across embryo and endosperm tissues.

### Slide 6
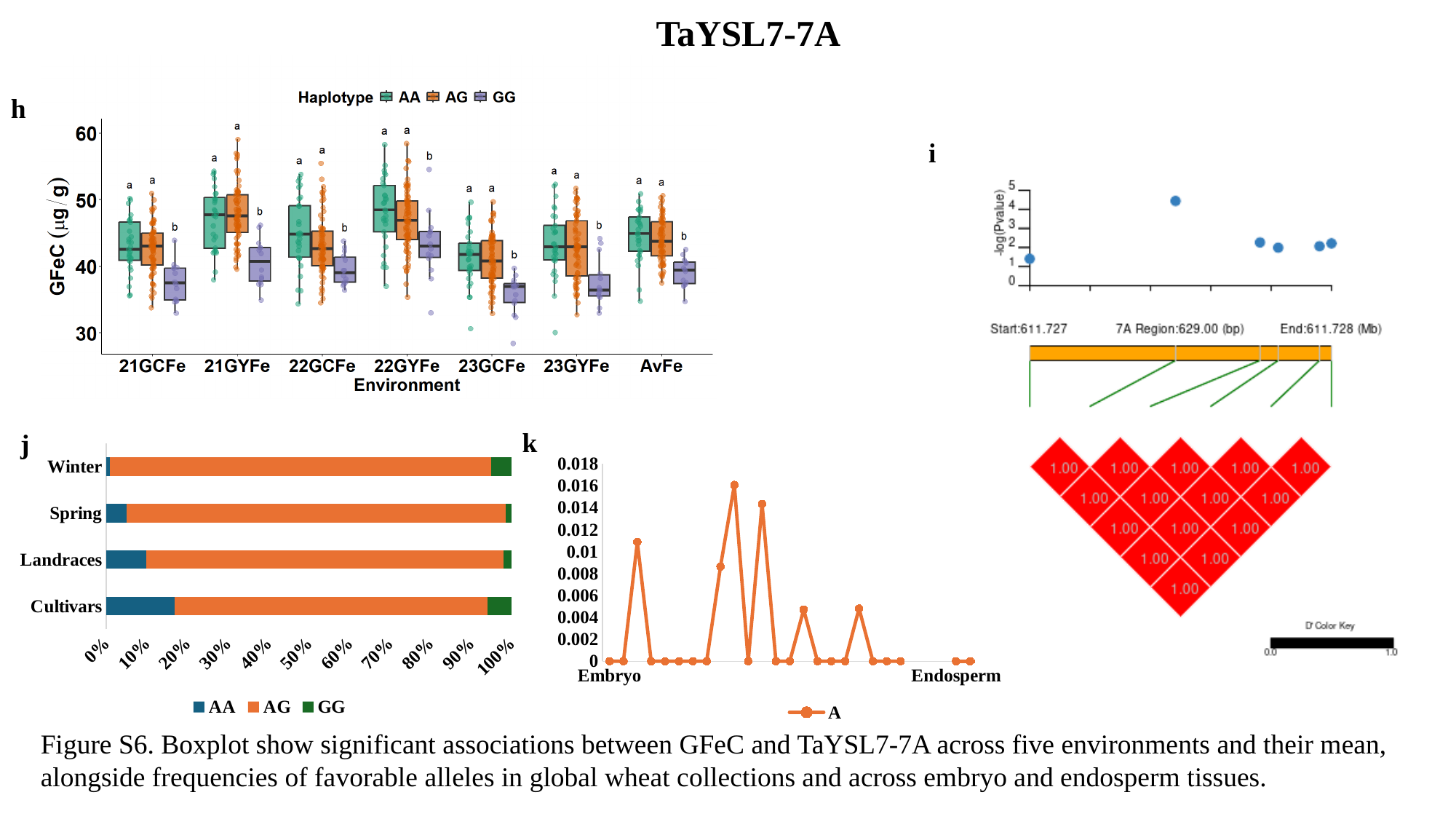

TaYSL7-7A
h
i
k
j
#### Chart
| Category | AA | AG | GG |
|---|---|---|---|
| Cultivars | 91.0 | 415.0 | 32.0 |
| Landraces | 62.0 | 552.0 | 13.0 |
| Spring | 7.0 | 133.0 | 2.0 |
| Winter | 7.0 | 770.0 | 41.0 |
#### Chart
| Category | A |
|---|---|
| Embryo | 0.0 |
| | 0.0 |
| | 0.0108667 |
| | 0.0 |
| | 0.0 |
| | 0.0 |
| | 0.0 |
| | 0.0 |
| | 0.00863333 |
| | 0.0160667 |
| | 0.0 |
| | 0.0143333 |
| | 0.0 |
| | 0.0 |
| | 0.0047 |
| | 0.0 |
| | 0.0 |
| | 0.0 |
| | 0.0048 |
| | 0.0 |
| | 0.0 |
| | 0.0 |
| | None |
| | None |
| | None |
| Endosperm | 0.0 |
| | 0.0 |
| | 0.0 |
| | 0.0 |
| | 0.0 |
| | 0.0 |
| | 0.0 |
| | 0.0 |
| | 0.0 |Figure S6. Boxplot show significant associations between GFeC and TaYSL7-7A across five environments and their mean, alongside frequencies of favorable alleles in global wheat collections and across embryo and endosperm tissues.
